# Lung-to-ear sound transmission does not improve directional hearing in green treefrogs (*Hyla cinerea*)

**DOI:** 10.1101/2020.06.30.171074

**Authors:** Jakob Christensen-Dalsgaard, Norman Lee, Mark A. Bee

## Abstract

Amphibians are unique among extant vertebrates in having middle ear cavities that are internally coupled to each other and to the lungs. In frogs, the lung-to-ear sound transmission pathway can influence the tympanum’s inherent directionality, but what role such effects might play in directional hearing remain unclear. In this study of the American green treefrog (*Hyla cinerea*), we tested the hypothesis that the lung-to-ear sound transmission pathway functions to improve directional hearing, particularly in the context of interspecific sexual communication. Using laser vibrometry, we measured the tympanum’s vibration amplitude in females in response to a frequency modulated sweep presented from 12 sound incidence angles in azimuth. Tympanum directionality was determined across three states of lung inflation (inflated, deflated, reinflated) both for a single tympanum in the form of the vibration amplitude difference (VAD) and for binaural comparisons in the form of the interaural vibration amplitude difference (IVAD). The state of lung inflation had negligible effects (typically less than 0.5 dB) on both VADs and IVADs at frequencies emphasized in the advertisement calls produced by conspecific males (834 Hz and 2730 Hz). Directionality at the peak resonance frequency of the lungs (1558 Hz) was improved by ≅ 3 dB for a single tympanum when the lungs were inflated versus deflated, but IVADs were not impacted by the state of lung inflation. Based on these results, we reject the hypothesis that the lung-to-ear sound transmission pathway functions to improve directional hearing in frogs.

**SUMMARY STATEMENT:** Contrary to prevailing views on the mechanisms of hearing in frogs, the lung-to-ear pathway for sound transmission does not improve directional hearing in these vociferous vertebrates.

## INTRODUCTION

Vocal communication is of prime importance for successful reproduction in most frogs (Gerhardt and Huber, 2002; Narins et al., 2007; Ryan, 2001). In many species, stationary male signalers produce loud advertisement calls to attract females to their calling site or territory (Schwartz and Bee, 2013; Wells and Schwartz, 2007). As an intended receiver, a gravid female must perform two critical perceptual tasks in order to reproduce (Gerhardt and Bee, 2007). First, she must recognize that a male’s call belongs to the set of calls of her species based on analyzing its spectral and temporal properties. Second, she must localize and approach the source of the calls using phonotaxis (approach toward sound), often while navigating a spatially complex microhabitat under low light levels, such as a vegetated wetland at night. Both perceptual tasks – sound pattern recognition and sound source localization – commonly take place in noisy breeding choruses that create unfavorable signal-to-noise ratios for communication (Bee, 2012, 2015; Bee and Christensen-Dalsgaard, 2016; Vélez et al., 2013).

Previous studies have shown that frogs exhibit highly directional phonotaxis toward conspecific calls, with jumps and head orientations often directed to within a few degrees of the sound source (Feng et al., 1976; Gerhardt and Rheinlaender, 1982; Jørgensen and Gerhardt, 1991; Rheinlaender et al., 1979; Shen et al., 2008; Ursprung et al., 2009). In addition, frogs have good source localization acuity and angle discrimination for sounds originating within the frontal field (e.g., ±45° from the midline)(Caldwell and Bee, 2014; Klump and Gerhardt, 1989). Although directional phonotaxis is a critically important component of frog reproductive behavior, we still lack a complete understanding of the mechanisms underlying directional hearing and sound source localization in these animals (Bee and Christensen-Dalsgaard, 2016; Eggermont, 1988; Klump, 1995; Rheinlaender and Klump, 1988).

Two anatomical features of the frog’s auditory system are currently believed to play key functional roles in directional hearing: their internally coupled ears and their lungs (Bee and Christensen-Dalsgaard, 2016). The free-field directionality of the frog’s tympanum has been investigated in a few species, in all cases showing a directional response at frequencies where diffraction of sound around the head is unimportant (Caldwell et al., 2014; Jørgensen, 1991; Jørgensen and Gerhardt, 1991; Jørgensen et al., 1991; Vlaming et al., 1984). Most likely, a part of the increased directionality is produced by acoustical interaction of the two tympana, because the air-filled middle ears of most frogs are internally coupled through the mouth cavity via wide and permanently open Eustachian tubes (Christensen-Dalsgaard, 2005, 2011; van Hemmen et al., 2016; but see Gridi-Papp et al., 2008). With this physical arrangement, each tympanum functions as a pressure-difference receiver with an inherently directional vibration response to sound originating from different positions in azimuth, although sound input to both ears is still required for accurate sound localization (Feng et al., 1976). Frogs also possess an accessory pathway for sound transmission between the body wall overlying the lungs and the internally-coupled middle ears (Ehret et al., 1990, 1994; Narins et al., 1988). The lungs can influence the magnitude of the tympanum’s directionality, with these effects being more pronounced at frequencies near the resonance frequency of the lungs (Jørgensen, 1991; Jørgensen et al., 1991; see also reviews in Christensen-Dalsgaard, 2005, 2011). While internally coupled ears also play functional roles in directional hearing in some other vertebrates, such as lizards and crocodilians (Bierman et al., 2014; Carr and Christensen-Dalsgaard, 2016; Christensen-Dalsgaard and Manley, 2008; Christensen-Dalsgaard et al., 2011), and in some invertebrates, such as crickets and katydids (Römer, 2015; Römer and Schmidt, 2016), the frog’s lung-to-ear sound transmission pathway appears to be unique among vertebrates, and its precise contribution to directional hearing remains uncertain (Bee and Christensen-Dalsgaard, 2016). At present, each tympanum’s directional response is thought to result from the interaction of sound impinging directly on its external surface and sound that indirectly reaches its internal surface from the other tympanum via the internal coupling and from the lungs via the glottis. Although there is strong empirical and theoretical support for the role of internal coupling in determining the tympanum’s directionality, and there is clear evidence that the lungs contribute to the tympanum’s response to sound, how these multiple inputs – direct from the free field, indirect via the opposite tympanum, and indirect via the lungs – act together to influence directional hearing in frogs is not well understood

In this study of the American green treefrog, *Hyla cinerea* (Schneider, 1799), we tested the hypothesis that the lung-to-ear sound transmission pathway functions to improve directional hearing, particularly in the context of interspecific sexual communication. The auditory and vocal communication systems of green treefrogs have been well studied in terms of basic auditory perception and physiology (Buerkle et al., 2014; Ehret and Capranica, 1980; Ehret and Gerhardt, 1980; Ehret et al., 1983; Gerhardt and Höbel, 2005; Klump et al., 2004; Megela-Simmons et al., 1985; Moss and Simmons, 1986; Penna et al., 1992); sound pattern recognition, species recognition, and mate choice (Allan and Simmons, 1994; Gerhardt, 1974, 1978a, b, 1981, 1987, 1991; Gerhardt et al., 1987; Höbel and Gerhardt, 2003; Lee et al., 2017; Simmons, 1988; Simmons et al., 1993); and sound source localization (Feng et al., 1976; Klump et al., 2004; Rheinlaender et al., 1979). In contrast to the extensive data on neurophysiology and behavior, only two studies have investigated the biophysics, and in particular the directionality, of the tympanum in green treefrogs (Michelsen et al., 1986; Rheinlaender et al., 1981). However, neither study investigated the role of the lung-to-ear sound transmission pathway in determining the directionality of the tympanum.

We used laser vibrometry to quantify the contribution of the lung-to-ear pathway in shaping the inherent directionality of the tympanum’s vibration amplitude response. To this end, we investigated how both the *vibration amplitude difference* (VAD) for a single tympanum and the *interaural vibration amplitude difference* (IVAD) varied as functions of lung inflation in response to free-field stimulation. The VAD refers to the difference between vibration amplitudes of the measured tympanum across different angles of sound incidence at a particular frequency. Thus, as a measure of directionality, the VAD represents the physical change in how a single tympanum responds to sounds presented from different angles in azimuth. As a complement to the VAD, the IVAD is a computed measure of binaural disparity that is more relevant to questions of directional hearing because it quantifies the difference between the vibration amplitudes of the left and right tympana when sound of a particular frequency originates from a particular sound incidence angle (Jørgensen et al., 1991). Hence, the IVAD estimates the difference in input through the auditory periphery on the left and right sides of the animal and, therefore, more closely approximates the information available to the auditory nervous system for computing the azimuthal direction of a sound source (Bee and Christensen-Dalsgaard, 2016).

## MATERIALS AND METHODS

### Animals

All animal procedures were approved by the Institutional Animal Care and Use Committee of the University of Minnesota (#1401-31258A) and complied with the NIH *Guide for the Care and Use of Laboratory Animals* (8^th^ Edition). Subjects were 25 adult female green treefrogs of unknown age (mean ± SD snout-to-vent length = 53.61 ± 3.50 mm, range = 47.7 to 59.2 mm; mean ± SD mass = 12.71 ± 2.92 g, range = 8.3 to 17.6 g; mean ± SD interaural distance = 14.08 ± 1.00 mm, range = 11.9 to 15.7 mm). They were collected in amplexus in July 2015 from ponds located at the East Texas Conservation Center in Jasper County, Texas, U.S.A. (30°56’46.15”N, 94°7’51.46”W). Animals were returned within 48 hours of collection to the laboratory at the University of Minnesota, where they were housed on a 12-hour light:dark cycle, provided with access to perches and refugia, fed a diet of vitamin-dusted crickets, and given ad libitum access to fresh water.

All laser measurements of an individual subject were made during a single recording session of less than two hours during which they were immobilized with an intramuscular injection of succinylcholine chloride (5 μg/g) into the thigh. We allowed subjects to regulate their own lung volume undisturbed over the 5-10 minutes during which the immobilizing agent took effect. After full immobilization was achieved, the state of lung inflation resembled that observed in unmanipulated frogs sitting in a natural posture based on visual inspection of lateral body wall extensions (see Caldwell et al., 2014). Hereafter, we refer to this state of lung inflation as the “inflated” condition. We also took laser measurements in two additional states of lung inflation. To create the “deflated” condition, we expressed the air from the animal’s lungs by gently depressing the lateral body wall while using the narrow end of a small, plastic pipette tip to hold the glottis open. For the “reinflated” condition, we attempted to return the lungs to their natural level of inflation observed prior to manual deflation by gently blowing air by mouth through a pipette with its tip located just above the closed glottis; the movement of air was sufficient to open the glottis and inflate the lungs. We have previously shown these procedures to be effective at deflating and reinflating the lungs (Caldwell et al., 2014; Lee et al., 2020). By necessity, measurements were always made first in the inflated condition followed by the deflated condition and then the reinflated condition. We facilitated cutaneous respiration while subjects were immobilized by periodically applying water to the dorsum to keep the skin moist. Animals that resumed buccal pumping during a recording session received a second, half dose of succinylcholine chloride. In the analyses reported below, four of 25 subjects were excluded (final *N* = 21) because we could not confidently confirm differences in their state of lung inflation across treatments based on visual inspection of lateral body wall extensions.

### Apparatus

Laser measurements were made inside a custom-built, semi-anechoic sound chamber (inside dimensions: 2.9 m [L] × 2.7 m [W] × 1.9 m [H], Industrial Acoustics Company, North Aurora, IL). The inside walls and ceiling of the chamber were lined with Sonex acoustic foam panels (Model VLW-60; Pinta Acoustic, Inc. Minneapolis, MN) to reduce reverberations. The chamber’s floor was covered in low-pile carpet. During recordings, subjects sat in a typical posture in the horizontal plane atop an acoustically transparent, cylindrical pedestal (30-cm tall, 7-cm diameter) made from wire mesh (0.9-mm diameter wire, 10.0-mm grid spacing). A raised arch of thin wire was used to support the tip of the subject’s mandible, such that its jaw was parallel to the ground. We mounted the base of the pedestal to a horizontal, 70-cm long piece of Unistrut^®^ (Unistrut, Harvey, IL) that extended to the center of the sound chamber from a mount on a vibration isolation table (Technical Manufacturing Corporation, Peabody, MA) positioned against an inside wall of the sound chamber. We covered the Unistrut and vibration isolation table with the same acoustic foam that lined the walls and ceiling of the chamber. In this configuration, the top of the pedestal was 120 cm above the floor of the sound chamber at its center. The laser vibrometer used for measurements (PDV-100, Polytech, Irvine, CA) was mounted at the same height on the vibration isolation table from which the subject pedestal was mounted and was positioned 70° to the animal’s right relative to the position of its snout (0°).

### Acoustic stimulation and laser measurements

We presented subjects with a frequency-modulated (FM) sweep (44.1 kHz, 16-bit) broadcast at a sound pressure level (SPL re 20 μPa, fast, C-weighted) of 85 dB from each of 12 sound incidence angles (0° to 330° in 30° steps) in azimuth. The direction in which the animal’s snout pointed was taken to be 0°. Thus, an angle of +90 corresponds to the animal’s right side, which was ipsilateral to the laser, and an angle of 270° (equivalent to −90°) corresponds to the animal’s left side, which was contralateral to the laser. The stimulus had a total duration of 195 ms, with linear onset and offset ramps of 10 ms. Over the 175-ms steady-state portion of its amplitude envelope, the stimulus increased linearly in frequency from 0.2 to 7.5 kHz. We amplified the stimulus (Crown XLS1000, Elkhart, IN) and broadcast it through a single speaker (Mod1, Orb Audio, New York, NY) that was positioned 50 cm away from the approximate center of a subject’s head (measured from the intersection of the midline and the interaural axis) as it sat on the pedestal. The speaker was suspended from the ceiling of the sound chamber by a rotating arm fashioned from steel pipe and covered in acoustic foam that allowed us to put the speaker at any azimuthal position relative to the subject. The speaker was at the same height above the chamber floor as the subject. We used a G.R.A.S. 40SC probe microphone (G.R.A.S. Sound & Vibration A/S, Holte, Denmark) to record stimuli at the position of the tympanum. The tip of a flexible probe tube was positioned approximately 2 mm from the edge of the subject’s right tympanum during recordings. We amplified the output of the probe microphone using an MP-1 microphone pre-amplifier (Sound Devices, Reedsburg, WI) and recorded its output using an external digital and analog data acquisition (DAQ) device (NI USB 6259, National Instruments, Austin, TX).

Just prior to making recordings, we calibrated the stimulus separately for each speaker position using a Brüel & Kjær Type 2250 sound level meter (Brüel & Kjær Sound & Vibration Measurement A/S, Nærum, Denmark) and a Brüel & Kjær Type 4189 ½-inch condenser microphone. Signal levels for calibration and playback were controlled using a programmable attenuator (PA5, Tucker-Davis Technologies, Alachua, FL). For calibration measurements, the microphone of the sound level meter was suspended from the ceiling of the sound chamber by an extension cable (AO-0414-D-100) so that the tip of the microphone was located at the position occupied by a subject’s head during recordings.

For each subject, the vibration amplitude of the right tympanum was first recorded in the inflated condition, beginning at a randomly determined location around the subject. Responses at each successive angle were recorded after repositioning the speaker in a counterclockwise direction. At each speaker location, we recorded the tympanum’s response to 20 repetitions of the stimulus (1.5-s stimulus period). After recording responses from the 12th and final speaker location in the inflated condition, we repeated the above procedure to measure the tympanum in the deflated condition beginning at the same randomly determined starting location used in the naturally inflated condition. Following all measurements in the deflated condition, this procedure was then repeated a third and final time in the reinflated condition.

We measured the vibration amplitude of the animal’s right tympanum by focusing the laser on a small (45-μm to 63-μm diameter), retroreflective glass bead (P-RETRO-500, Polytech, Irvine, CA) placed at the center of the right tympanum. We digitized (44.1 kHz, 16 bit) the analog output of the laser using the NIDAQ device, which we controlled using MATLAB^®^ (v.2014a, MathWorks, Natick, MA) running on an OptiPlex 745 PC (Dell, Round Rock, TX). Tympanum vibration spectra were calculated from the acquired laser signals in MATLAB using the pwelch function (window size = 256, overlap = 50%). Vibration spectra were corrected for small directional variation in the sound spectrum by calculating transfer functions between tympanum vibrations and sound at the tympanum’s external surface. This was done by dividing the tympanum vibration spectrum by the sound spectrum recorded by the probe microphone (i.e., by subtraction of dB values).

### Data analysis

We computed the VAD at a particular frequency as the maximum difference in the vibration amplitudes (in dB) of the measured tympanum that occurred across different angles of sound incidence. Computed this way, the VAD is a measure of the directional response of a single tympanum. In contrast, the IVAD, which is a measure of binaural disparity, was computed for a particular frequency from measurements of a single ear by assuming bilateral symmetry and computing the difference based on the correct corresponding angles. For example, the IVAD for 30° was computed as the difference between the vibration amplitudes measured at 30° and 330° (i.e., −30°) relative to the midline at 0°. VADs and IVADs were computed based on laser measurements made while the animals’ lungs were inflated and again after both deflating and reinflating the lungs. We restricted analyses of VADs and IVADs to those measured across a 4-octave range between 300 Hz and 4800 Hz to avoid artificially large values that were sometimes observed to occur well outside the range of typical tympanum sensitivity when the laser signal was near its internal noise floor. We used linear interpolation to determine the values of VADs and IVADs occurring between fixed angles of stimulation.

We focused our statistical analyses of VADs and IVADs on three frequencies of biological relevance: two spectral peaks emphasized in the advertisement calls produced by conspecific males and the peak resonance frequency of the lungs of females. Similar to related species in the genus *Hyla* (Gerhardt, 2001, 2005; Gerhardt et al., 2007), male green treefrogs produce an advertisement call with a frequency spectrum consisting of two prominent spectral peaks that are analogous to the formant frequencies present in human vowel sounds (Lee et al., 2017; Oldham and Gerhardt, 1975). The lower spectral peak is important for long-distance communication and source localization (Gerhardt, 1976; Klump et al., 2004; Rheinlaender et al., 1979), whereas the higher spectral peak may be more important in sexual selection via female mate choice (Gerhardt, 1976, 1981). Females have robust preferences for calls containing both spectral peaks (Lee et al., 2017). Based on analyses of a sample of 457 advertisement calls (≅ 20 calls from each of 23 males), Lee et al. (2020) reported the mean (± SD) frequencies of the lower and higher spectral peaks to be 834 ± 14 Hz and 2730 ± 34 kHz, respectively. The mean frequency of the “valley” between these two spectral peaks was 1653 ± 39 Hz. Using laser vibrometry, Lee et al. (2020) determined the mean peak frequency of the lung resonance in a sample of 10 females to be 1558 Hz (range: 1400 Hz to 1850 Hz).

A two-way repeated-measures analysis of variance (rmANOVA) was used to assess differences in the magnitude of the VAD as a function of lung inflation (inflated, deflated, and reinflated) at three fixed frequencies (834 Hz, 1558 Hz, and 2730 Hz). We used a three-way rmANOVA to assess differences in the magnitude of the IVAD as a function of lung inflation (inflated, deflated, and reinflated) at the same three fixed frequencies (834 Hz, 1558 Hz, and 2730 Hz) and at five fixed sound incidence angles (30° to 150° in 30° steps). We additionally report the results from quadratic contrasts for all main effects of lung inflation given the general expectation based on previous studies that the directionality observed with the lungs inflated should decrease in the deflated condition and be restored in the reinflated condition. For all rmANOVAs, we report F values (and their unadjusted degrees of freedom) based on the Greenhouse and Geiser (1959) correction for possible violations of sphericity and partial eta^2^ values as measures of effect size. We used α = 0.05 to determine statistical significance. All statistical analyses were conducted using SPSS v21.

## RESULTS

In all three lung inflation conditions, the tympanum had a bandpass frequency response and was most sensitive to frequencies between about 1000 Hz and 5000 Hz across most angles of sound incidence (Fig. 1). In both the inflated (Fig. 1A) and reinflated (Fig. 1C) conditions, there was also a prominent “dip” in the vibration amplitude spectrum between approximately 1400 Hz and 2200 Hz. This dip corresponded closely with the peak resonance frequency of the lungs (1558 Hz) and was largely absent in the deflated condition (Fig. 1B). Across frequencies, vibration amplitudes were greatest at ipsilateral angles of +60° to +120° (on the animal’s right side) compared with the corresponding contralateral angles of −60° to −120° (on the animal’s left side). The tympanum’s greatest directionality, as indicated by the differences in vibration amplitude between ipsilateral and contralateral angles, occurred in the frequency region of the dip (Fig. 1). These general trends are evident both in the tympanum’s vibration amplitude response for two representative frogs and in the mean vibration amplitude of the right tympanum averaged across all individuals (Fig. 1). Reinflation of the deflated lungs was more successful in some animals than in others at restoring the dip in the vibration amplitude spectrum (cf. Frog 1 and Frog 2 in Fig. 1C). Consequently, the directionality associated with the dip was, on average, slightly less pronounced in the reinflated condition compared with the inflated condition. We cannot rule out the possibility that changes in the volume of air in the mouth cavity resulting from manipulations of lung inflation also contribute to differences between the inflated and reinflated states.

**Fig. 1.**
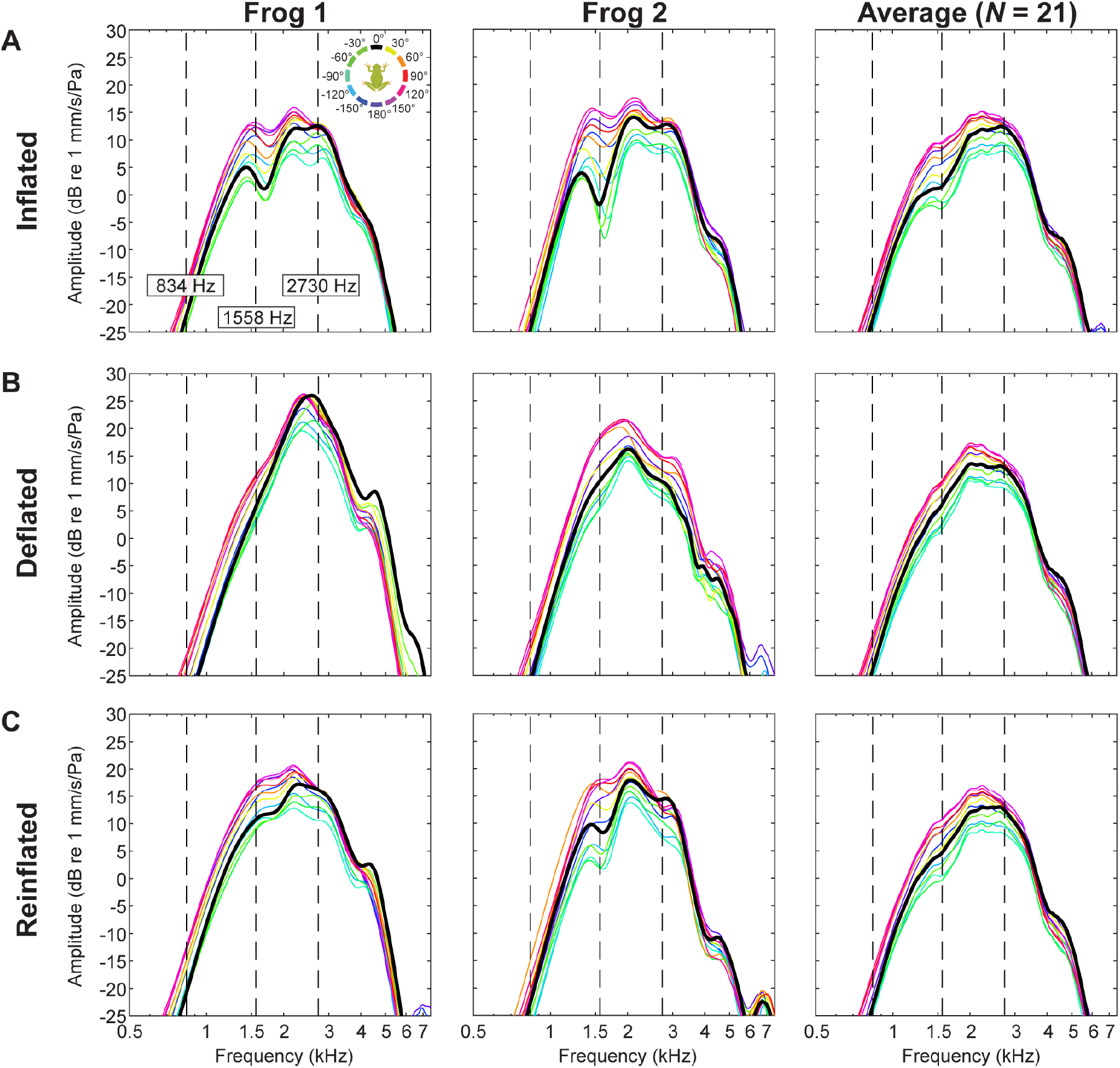
Response of the tympanum to free-field sound in three states of lung inflation. Shown here are tympanum transfer functions measured in response to a frequency modulated (FM) sweep broadcast from 12 sound incidence angles in azimuth (0, ±30°, ±60°, ±90°, ±120°, ±150°, 180°). Transfer functions at each angle were generated by dividing the vibration amplitude spectrum measured with the laser by the corresponding sound spectrum measured with the probe microphone adjacent to the measured tympanum. Data are presented for two representative frogs and averaged over 21 frogs and are shown separately for the (A) inflated, (B), deflated, and (C) reinflated states of lung inflation. Vertical dashed lines depict frequencies of the two spectral peaks in conspecific advertisement calls (834 Hz and 2730 Hz) and the peak resonance frequency of the lungs (1559 Hz) (Lee et al., 2020).

### Vibration amplitude differences (VADs)

With inflated lungs, the tympanum’s maximum directionality – as measured by the maximum VAD at any frequency across any two sound incidence angles – was 15.5 dB and occurred at a frequency of 1486 Hz and between angles of +99° (ipsilateral) and −49° (contralateral) (Table 1). The maximum VAD decreased by 4 dB when the lungs were deflated and was largely restored (within 1 dB) when the lungs were reinflated (Table 1). Differences in the mean frequency and mean sound incidence angles at which the maximum VAD occurred were generally small (e.g., < 240 Hz and < 40°) across the three states of lung inflation (Table 1).

**Table 1.**
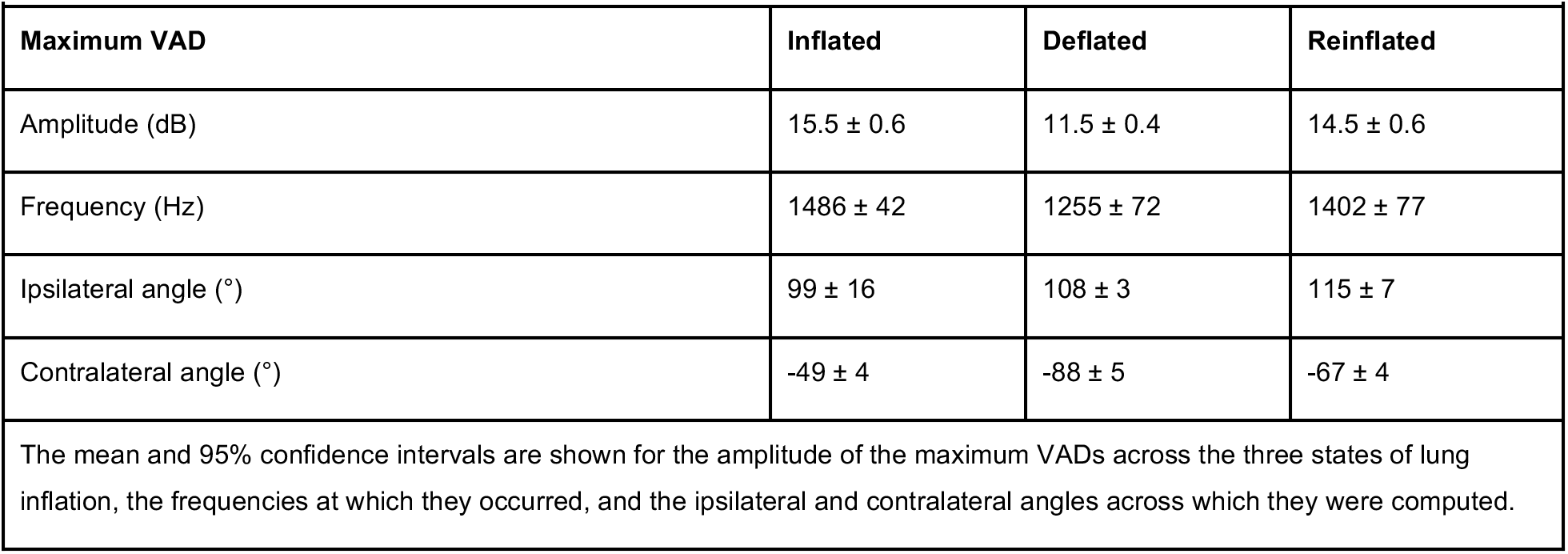
Summary of maximum vibration amplitude differences (VADs) in three states of lung inflation

We assessed the impacts of lung inflation on VADs using a 3 lung inflation (inflated, deflated, reinflated) × 3 frequency (834 Hz, 1558 Hz, 2730 Hz) rmANOVA. Recall that the lowest (834 Hz) and highest (2730 Hz) frequencies correspond to the two prominent spectral peaks in conspecific advertisement calls, and the intermediate frequency (1558 Hz) corresponds to the peak resonance frequency of the inflated lungs. This analysis revealed significant main effects of both lung inflation (F_2,40_ = 6.18, P = 0.006, partial eta^2^ = 0.24) and frequency (F_2,40_ = 52.98, P < 0.001, partial eta^2^ = 0.73), and their two-way interaction was also significant (F_4,80_ = 5.55, P = 0.002, partial eta^2^ = 0.22). The quadratic contrast for the main effect of lung inflation was also significant (F_1,20_ = 14.94, P = 0.001, partial eta^2^ = 0.43). Averaged across all three frequencies, the mean (± 95% CI) VADs were lowest in the deflated condition (8.5 ± 0.7 dB) and higher and more similar in the inflated (9.8 ± 0.7 dB) and reinflated (10.1 ± 1.1 dB) conditions. Averaged across all three states of lung inflation, mean (± 95% CI) VADs were largest at a frequency of 1558 Hz (11.6 ± 1.1 dB), smallest at 2730 Hz (6.8 ± 0.4 dB), and intermediate at 834 Hz (10.1 ± 0.9 dB). The two-way interaction between lung inflation and frequency resulted because the impacts of lung inflation were more pronounced at the intermediate frequency of 1558 Hz compared with call frequencies of 834 Hz and 2730 Hz (Fig. 2). At 1558 Hz, the mean VAD was 3.4 dB and 2.6 dB lower in the deflated condition compared with the inflated and reinflated conditions, respectively (Fig. 2). In contrast, the differences in mean VADs across the three states of lung inflation were less than 1.8 dB, and most were less than 0.4 dB, at frequencies of 834 Hz and 2730 Hz (Fig. 2).

**Fig. 2.**
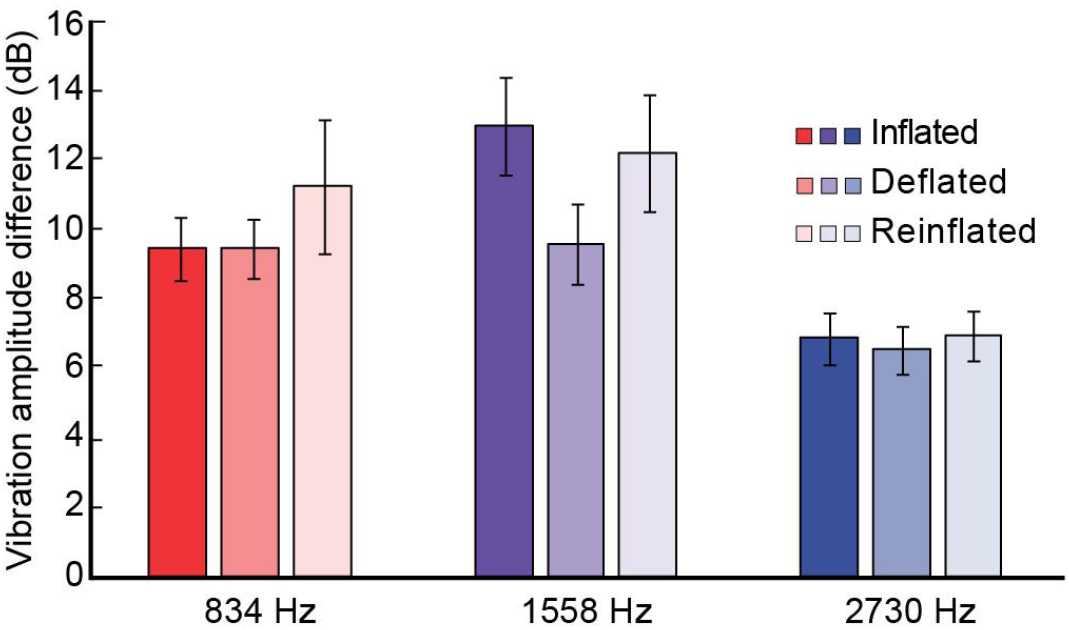
Impacts of lung inflation on the vibration amplitude difference (VAD). Shown here are the mean (± 95% CIs) VADs measured in three states of lung inflation (inflated, deflated, and reinflated) at frequencies corresponding to the two spectral components emphasized in conspecific calls (834 Hz and 2730 Hz) and at the peak resonance frequency of the lungs (1558 Hz).

### Interaural vibration amplitude differences (IVADs)

The left-right asymmetries present in measures of vibration amplitude (Fig. 1) are also reflected in the magnitudes of the IVADs plotted as heatmaps in Figure 3. IVADs in all states of lung inflation were generally greatest between about 1000 Hz and 3000 kHz and between +45° and +135°, with the largest IVADs occurring near 1600 Hz at 90° (Figure 3). Figure 4 depicts IVADs as functions of lung inflation at the two call frequencies and the lung resonance frequency, and Figure 5 illustrates how these IVADs varied as a function of sound incidence angle. We evaluated how IVADs varied as a function of lung inflation at fixed combinations of sound incidence angle and frequency using a 3 lung inflation (inflated, deflated, reinflated) × 5 angle (30°, 60°, 90°, 120°, 150°) × 3 frequency (834 Hz, 1558 Hz, 2730 Hz) rmANOVA. There were large and significant main effects of angle (F_4,80_ = 113.2, P < 0.001, partial eta^2^ = 0.87) and frequency (F_2,40_ = 41.5, P < 0.001, partial eta^2^ = 0.68), and there was a significant two-way interaction between angle and frequency (F_8,160_ = 26.8, P < 0.001, partial eta^2^ = 0.57). As illustrated in the polar plots in Figure 5, IVADs tended to be highest near sound incidence angles close to 90° and to decrease to 0 dB as the sound source was positioned closer to the axis of the midline (0° and 180°). Mean (± 95% CI) IVADs, averaged across sound incidence angles and states of lung inflation, were higher at the peak resonance frequency of the lungs (1558 Hz: 6.6 ± 1.6 dB) and lower at the two frequencies emphasized in conspecific mating calls (834 Hz: 5.2 ± 1.2 dB; 2730 Hz: 3.5 ± 0.7) (Figs. 4 & 5). The largest IVAD occurred at 90° and 1558 Hz, and the smallest IVAD occurred at 30° and 2730 Hz (Fig. 5).

**Fig. 3.**
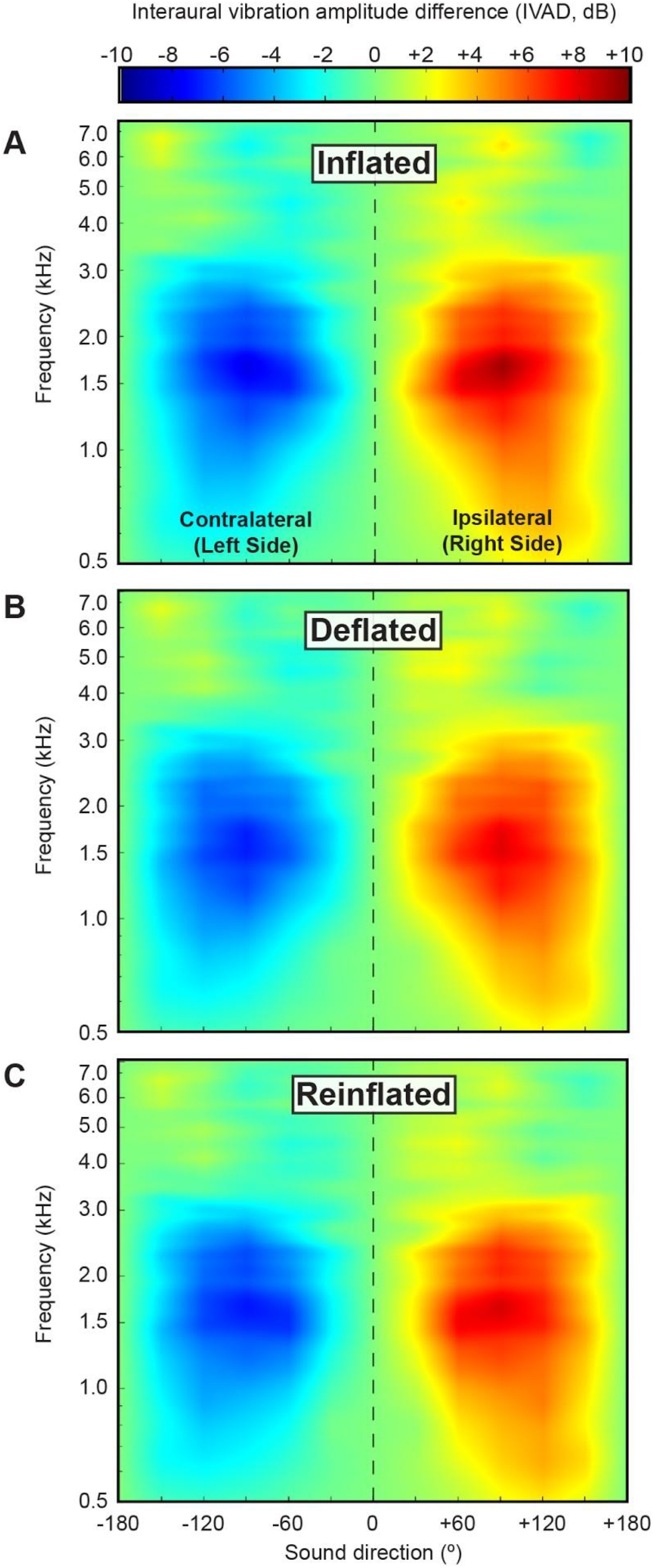
Interaural vibration amplitude differences (IVADs). Shown here are the mean interaural vibration amplitude differences (IVADs; *N* = 21) in response to a frequency-modulated (FM) sweep as a function of frequency and sound incidence angle. In each plot, values are interpolated across frequency and sound incidence angle and averaged over 21 subjects. Data are shown separately for the (A) inflated, (B) deflated, and (C) reinflated states of lung inflation. IVADs are measures of binaural disparity that assume bilateral symmetry. These heatmap plots of IVADs depict the differential stimulation of the ipsilateral (right) side, with positive values (red color) corresponding to sounds arising in the ipsilateral hemifield and negative values (blue color) corresponding to sounds arising in the contralateral hemifield.

**Fig. 4.**
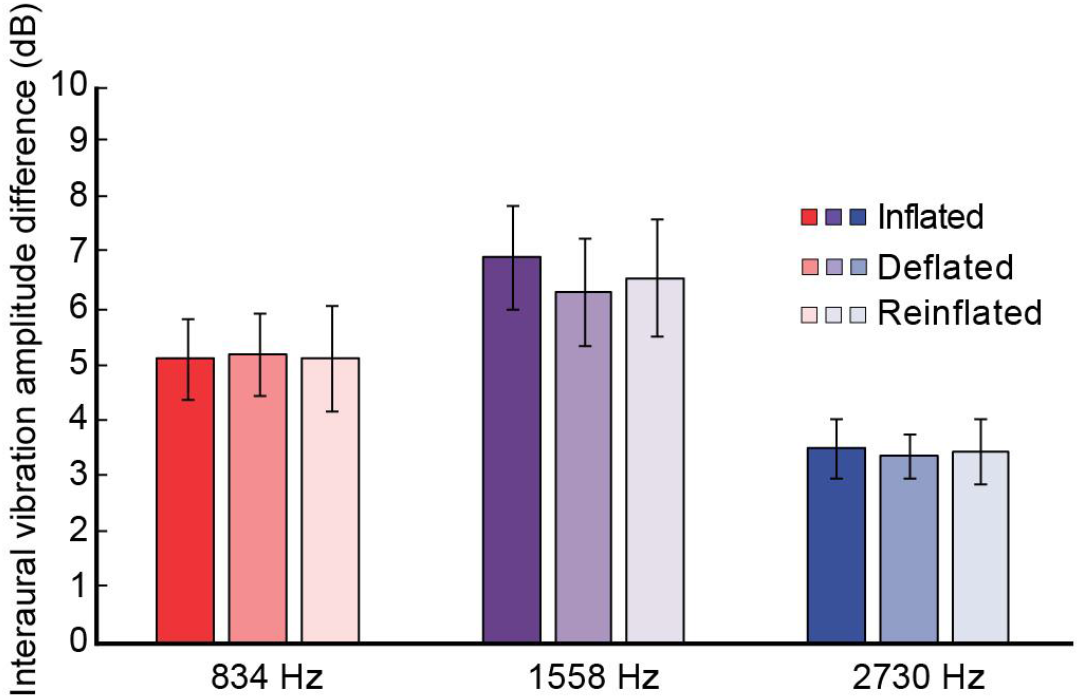
Impacts of lung inflation on the interaural vibration amplitude difference (IVAD). Shown here are the mean (± 95% CIs) IVADs measured in three states of lung inflation (inflated, deflated, and reinflated) at frequencies corresponding to the two spectral components emphasized in conspecific calls (834 Hz and 2730 Hz) and at the peak resonance frequency of the lungs (1558 Hz). Values are averaged over the measured sound incidence angles of 30°, 60°, 90°, 120°, and 150°.

**Fig. 5.**
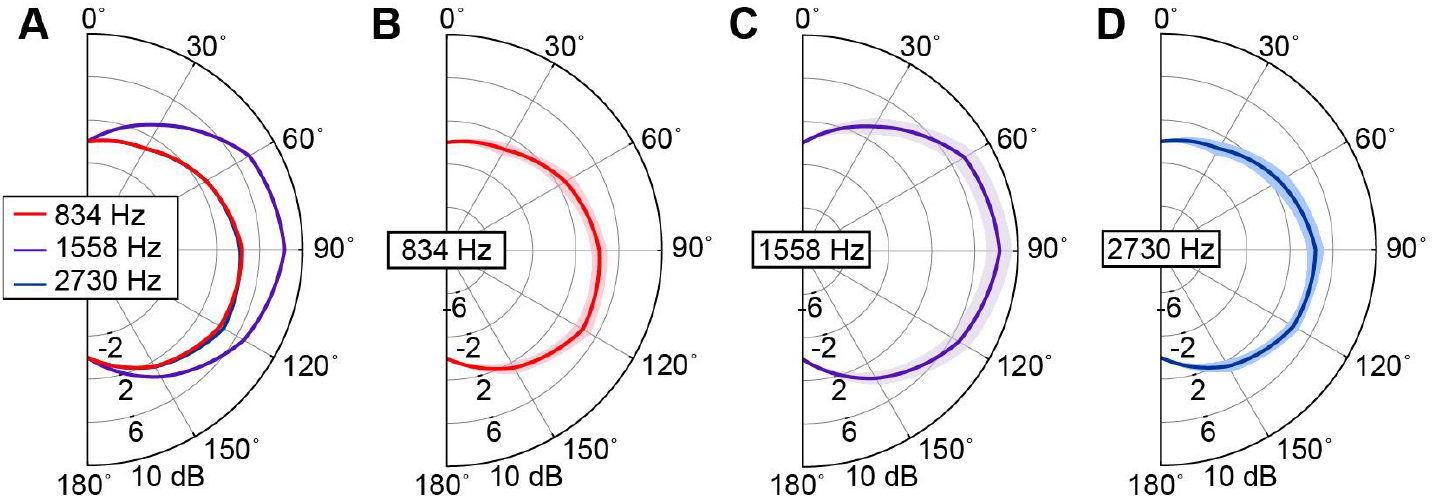
Interaural vibration amplitude differences (IVADs) as functions of frequency and sound incidence angle. (A) Half-polar plot comparing the magnitude of the mean IVAD in the inflated condition as a function of sound incidence angle for two frequencies emphasized in conspecific advertisement calls (834 Hz and 2730 Hz) and at the mean peak frequency of the lung resonance (1558 Hz). IVADs were greatest at all three frequencies when sounds were presented from lateral positions. (B-C) Half polar plots showing the mean (line) ± 95% confidence interval (shaded region) IVAD separately for each frequency as a function of sound incidence angle.

In contrast to the effects of sound incidence angle and frequency, the state of lung inflation had no significant effects on IVADs (Fig. 4): both the main effect of lung inflation (F_2,40_ = 0.37, P = 0.680, partial eta^2^ = 0.02) and the quadratic contrast for the main effect of lung inflation (F_1,20_ = 0.48, P = 0.495, partial eta^2^ = 0.02) were small and nonsignificant, as were the two-way interactions between lung inflation and both angle (F_8,160_ = 1,40, P = 0.245, partial eta^2^ = 0.07) and frequency (F_4,80_ = 0.45, P = 0.710, partial eta^2^ = 0.02), as well as the three-way interaction between these variables (F_16,320_ = 0.96, P = 0.53, partial eta^2^ = 0.05). The overall mean (± 95% CIs) IVADs, averaged over angle and frequency, were 5.2 ± 1.1 dB, 5.0 ± 1.2 dB, and 5.1 ± 1.1 dB in the inflated, deflated, and reinflated conditions.

The difference in IVADs between the inflated and deflated states of lung inflation were close to 0 dB at all combinations of frequency and sound incidence angle. This outcome is apparent in the heatmap shown in Figure 6A, which illustrates the mean differences in IVADs between the inflated and deflated states across sound incidence angles and frequency in relation to the average spectrum of conspecific calls. The nearly uniform, pale green color of the heatmap corresponds to an IVAD near 0 dB and indicates that the state of lung inflation had a negligible impact on IVAD magnitude. At the two spectral peaks emphasized in conspecific calls, and at the peak resonance frequency of the lungs, the mean (± 95% CIs) IVAD differences between the inflated and deflated conditions included 0 dB at nearly all angles of sound incidence (Fig. 6B).

**Fig. 6.**
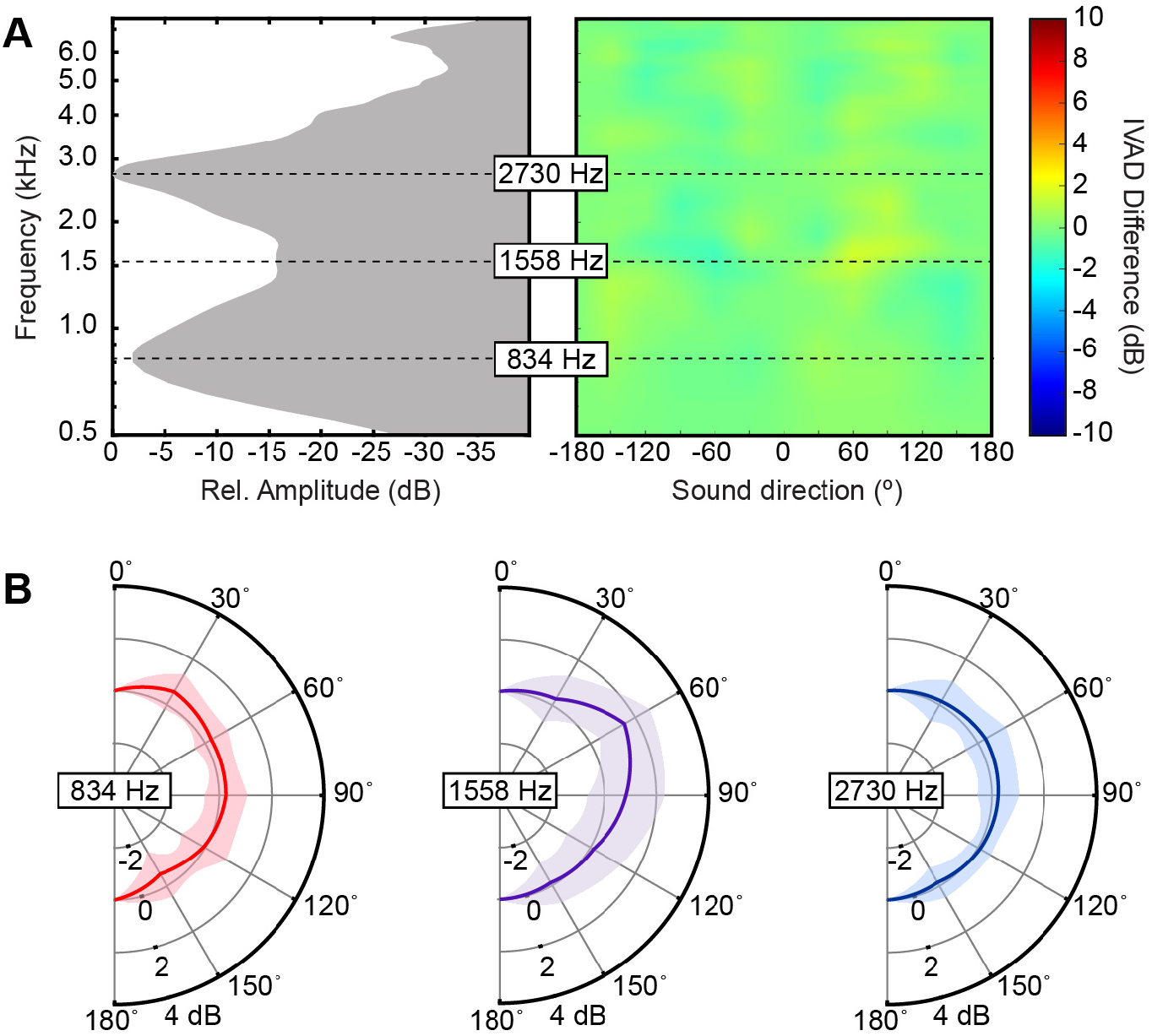
Relationship between the advertisement call spectrum and lung mediated differences in interaural vibration amplitude differences (IVADs). (A) On the left is shown the frequency spectrum of conspecific advertisement calls depicting relative amplitude as a function of frequency (redrawn from Lee et al., 2020). On the right is shown a heatmap of the differences in IVADs between the inflated and deflated conditions (inflated - deflated). Note that the call spectrum is rotated 90° counterclockwise relative to customary depictions of call spectra in order to facilitate comparisons with lung mediated differences in IVADs. The mean spectral peaks in the call (834 Hz and 2730 Hz) and the mean peak frequency of the lung resonance are highlighted. (B) Half polar plots depicting the magnitude (mean ± 95% CI) of the difference in IVADs between the inflated and deflated conditions as a function of sound incidence angle for the three frequencies highlighted in (A).

Table 2 summarizes the magnitudes of the maximum IVAD in each lung inflation condition along with the frequencies and sound incidence angles at which they occurred. On average, the magnitude of the maximum IVAD was similar in the inflated (12.4 dB; Table 2) and reinflated (12.5 dB; Table 2) conditions, and it was approximately 1 dB less in the deflated condition (11.2 dB; Table 2). Across the three lung inflation conditions, the mean frequencies at which the maximum IVAD was observed varied over a narrow range between 1579 Hz and 1690 Hz (Table 2), which is close to the mean peak resonance frequency of the lungs (1558 Hz). The mean angles at which the maximum IVAD was observed varied between 82.9° in the inflated condition and 100.0° in the deflated condition, and the median angle for all three states of lung inflation was 90° (Table 2).

**Table 2.** Summary of the maximum interaural vibration amplitude differences (IVADs) in three states of lung inflation

|  |  |  |  |  |  |  |  |  |  |  |  |  |
| --- | --- | --- | --- | --- | --- | --- | --- | --- | --- | --- | --- | --- |
|  | Magnitude of maximum IVAD (dB) |  |  |  | Frequency of maximum IVAD (Hz) |  |  |  | Angle of maximum IVAD (°) |  |  |  |
| Lung condition | Median | Mean | SD | CI | Median | Mean | SD | CI | Median | Mean | SD | CI |
| Inflated | 12.1 | 12.4 | 2.7 | 1.1 | 1550 | 1607.5 | 792.9 | 339.0 | 90 | 82.9 | 18.7 | 8.0 |
| Deflated | 10.6 | 11.2 | 2.4 | 1.0 | 1250 | 1689.9 | 1639.1 | 701.0 | 90 | 100.0 | 19.7 | 8.4 |
| Reinflated | 12.1 | 12.5 | 2.4 | 1.0 | 1450 | 1579.0 | 1439.7 | 615.8 | 90 | 95.7 | 22.5 | 9.6 |
| The median, mean, standard deviation (SD), and 95% confidence intervals (CI) are shown for the magnitude of the maximum IVAD, and the frequency and sound incidence angle at which the maximum occurred, based on a sample of 21 subjects. |  |  |  |  |  |  |  |  |  |  |  |  |

## DISCUSSION

The goal of this study was to test the hypothesis that the lung-to-ear sound transmission pathway functions to improve directional hearing. We found little evidence to support this hypothesis, and no evidence that sound input through the lungs might function to improve the localization of conspecific advertisement calls. Both the directionality of a single tympanum, as measured by the VAD, and the impacts of lung inflation on this measure of directionality were greater at frequencies near the peak resonance frequency of the lungs than at frequencies emphasized in conspecific calls. When measured as binaural disparities in the form of IVADs, directionality was again highest at frequencies near the peak resonance frequency of the lungs, but the state of lung inflation had little impact on directionality at these frequencies. Most importantly, the state of lung inflation had a negligible impact on directionality, measured either for a single ear (VADs) or as binaural disparities (IVADs), at the frequencies emphasized in conspecific calls. Based on these measurements, we conclude that the lung-to-ear sound transmission pathway probably plays no significant role in directional hearing in the context of localizing conspecific advertisement calls in green treefrogs.

Before discussing the role of the frog’s lungs in directional hearing in more detail, it is first worth considering how our findings on tympanum directionality relate to previous work in green treefrogs and other frog species. In their study of green treefrogs, Feng et al. (1976) were the first to demonstrate that female frogs rely on binaural cues to localize calling males. Sound localization was completely disrupted when one tympanum was covered with a thin layer of silicone grease, with frogs repeatedly turning in the direction of their unaltered ear (see also Rheinlaender et al., 1979). Based on measures of jump angles during phonotaxis in closed-loop tests, Rheinlaender et al. (1979; see also Klump et al., 2004) showed that sound localization in green treefrogs was highly accurate in azimuth, with a mean jump angle (i.e., the angle between the direction of the speaker and the direction of a jump) of 16.1° and a modal jump angle falling between 3° to 7°. Angles of head orientation (i.e., the angle between the direction of the speaker and the direction in which the frog’s snout pointed) were even smaller (mean = 8.4°; mode = 0° to 3°), indicating even better directional resolution than that based on jumps. Lateral head scanning prior to jumps improved jump accuracy in azimuth (mean jump angle = 11.8°; Rheinlaender et al., 1979), and scanning with the head raised above the horizontal plane was crucial for localizing elevated sources (Gerhardt and Rheinlaender, 1982). Klump and Gerhardt (1989) showed in open loop tests that the closely related barking treefrog (*Hyla gratiosa*) was capable of true angle discrimination, not just lateralization, and could discriminate angles differing by as little as 15° in azimuth within the frontal sound field (±45°), a result later corroborated by Caldwell and Bee (2014) for Cope’s gray treefrog (*Hyla chrysoscelis*). Rheinlaender et al. (1981) reported a directional sensitivity at 90° of about 4 dB in green treefrogs based on midbrain neural responses to a 1000 Hz tone. Using laser vibrometry to measure the tympanum’s response directly, Michelsen et al. (1986) later reported an mean IVAD at 90° closer to 9 dB in response to tones between 1000 Hz and 3000 Hz. These values generally accord well with the IVADs reported in the present study: at the call frequencies of 834 Hz and 2730 Hz, the mean (± 95% CI) IVADs at 90° were 4.4 ± 0.6 dB and 4.4 ± 0.7 dB, respectively, while that at the lung resonance of 1558 Hz was 8.9 ± 1.0 dB (Fig. 5). By comparison, sound pressure levels measured at the external surfaces of the tympana were typically ±1 dB in the range of 1000 Hz to 3000 Hz (Michelsen et al., 1986). Because we found the state of lung inflation had no impact on the magnitudes of IVADs, the larger binaural disparities in the tympanum’s response most likely arise from interaural coupling of the middle ears. In an electrophysiological study of sound localization in green treefrogs, Feng and Capranica (1978) reported that about 42% of cells in the superior olivary nucleus and 88% of cells in the inferior colliculus (torus semicircularis) were sensitive to binaural input, with the vast majority of binaural cells exhibiting EI responses in which the cells were excited by contralateral stimulation and inhibited by ipsilateral stimulation. Some EI neurons in the inferior colliculus were sensitive to small binaural disparities of just 1 dB to 2 dB, suggesting that IVADs on the order of 4 dB to 8 dB would be more than sufficient to drive directional neural responses subserving sound localization behavior in this species. Interaural time differences of up to 1.3 ms in auditory nerve responses to the amplitude modulation in natural calls may produce additional cues for sound localization (Klump et al., 2004).

As illustrated in Table 3, the present study also corroborates four broad patterns reported in previous biophysical studies of tympanum directionality in both closely related congeneric species as well as more distantly related species. First, the frog tympanum exhibits a bandpass frequency response with a characteristic “dip” in vibration amplitude at intermediate frequencies (Caldwell et al., 2014; Chung et al., 1981; Ho and Narins, 2006; Jørgensen, 1991; Jørgensen and Gerhardt, 1991; Jørgensen et al., 1991; Pinder and Palmer, 1983; Wilczynski et al., 1987). Second, the dip in tympanum sensitivity occurs at frequencies that are not emphasized in the spectra of conspecific advertisement calls (Caldwell et al., 2014; Jørgensen, 1991; Jørgensen and Gerhardt, 1991; Jørgensen et al., 1991). Third, the dip typically coincides with the frequencies of maximum tympanum directionality (Jørgensen, 1991; Jørgensen and Gerhardt, 1991; Jørgensen et al., 1991). And finally, the frequency range of the dip corresponds closely to the peak resonance frequency of the lungs (Caldwell et al., 2014; Jørgensen, 1991; Jørgensen et al., 1991). Green treefrogs exemplify all four patterns. The tympanum was most responsive in the frequency range of about 1000 Hz to 5000 Hz, and the tympanum’s transfer function had a prominent dip between approximately 1400 Hz and 2200 Hz when the lungs were inflated. The dip frequency range fell between the lower (834 Hz) and upper (2730 Hz) spectral peak of the advertisement call and included the peak resonance frequency of the lungs (1558 Hz; range: 1400 Hz to 1850 Hz; Lee et al., 2020). The maximum VAD was 15.5 dB and occurred at 1486 Hz, which is between the two spectral peaks of the call and within the frequency range of the dip in the tympanum’s transfer function. By comparison, VADs were smaller (approximately 5 to 10 dB) at the frequencies emphasized in conspecific advertisement calls. These patterns are similar to those documented in other species studied to date (Table 3), including eastern gray treefrogs (*Hyla versicolor*, Hylidae), Cope’s gray treefrogs (*H. chrysoscelis*, Hylidae), barking treefrogs (*H. gratiosa*, Hylidae), coqui frogs (*Eleutherodactylus coqui*, Eleutherodactylidae), and grass frogs (*Rana temporaria*, Ranidae). The three taxonomic families represented in this pool of species are members of two superfamilies, Hyloidea (Hylidae and Eleutherodactylidae) and Ranoidea (Ranidae), that last shared a common ancestor some 155 million years ago (Kumar et al., 2017). Thus, the patterns observed in the frog species studied to date may be both ancient and taxonomically widespread, and perhaps characteristic of the more than 5000 species of frog in the suborder Neobatrachia within the Anura. Species in other suborders have not been investigated.

**Table 3.**
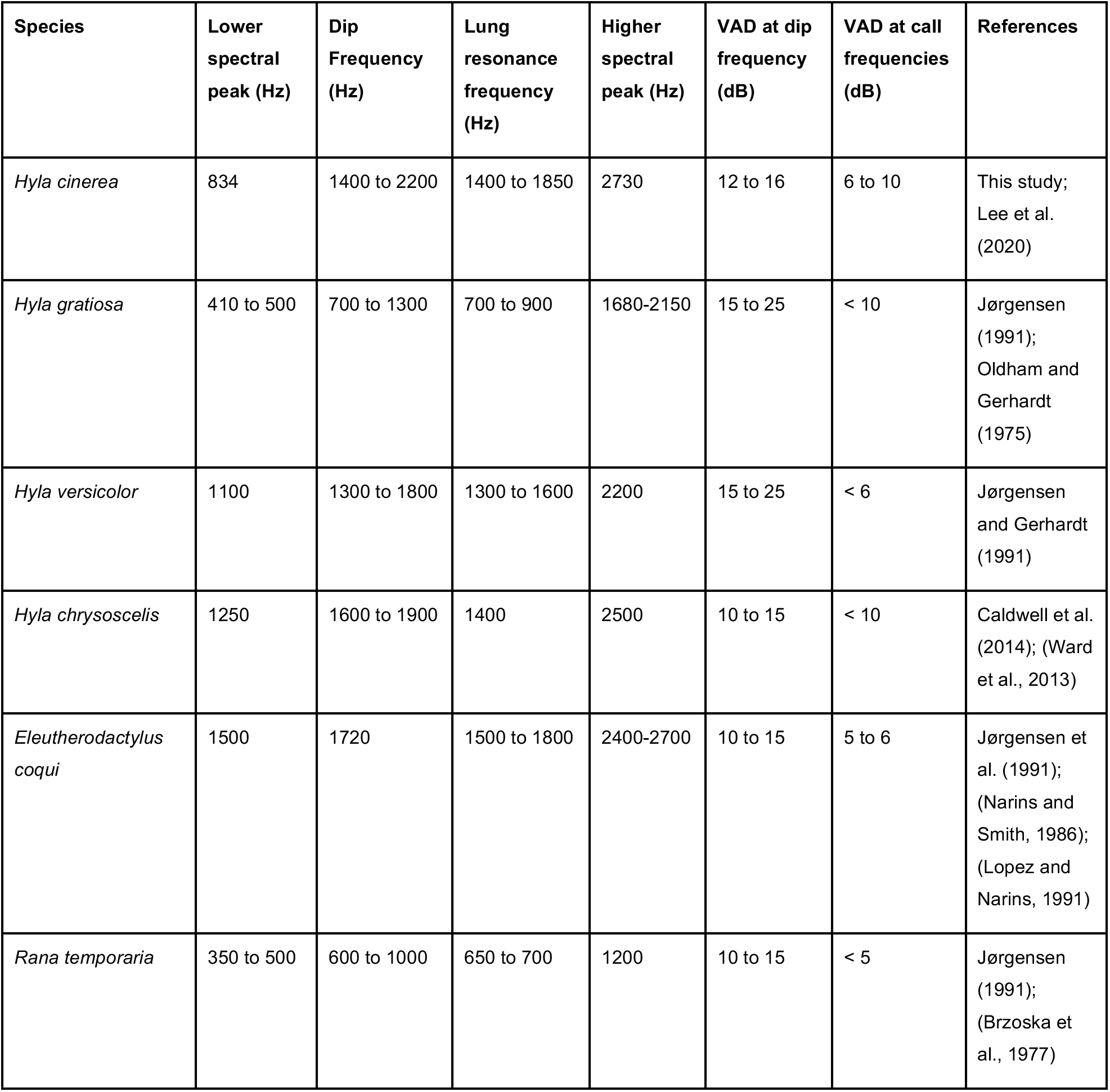
Relationships between frequencies of the “dip” in tympanum transfer functions, the spectral peaks of conspecific calls, the peak resonance of the lungs, and vibration amplitude differences (VADs)

|  |  |  |  |  |  |  |  |
| --- | --- | --- | --- | --- | --- | --- | --- |
| Species | Lower spectral peak (Hz) | Dip Frequency (Hz) | Lung resonance frequency (Hz) | Higher spectral peak (Hz) | VAD at dip frequency (dB) | VAD at call frequencies (dB) | References |
| *Hyla cinerea* | 834 | 1400 to 2200 | 1400 to 1850 | 2730 | 12 to 16 | 6 to 10 | This study; Lee et al. (2020) |
| *Hyla gratiosa* | 410 to 500 | 700 to 1300 | 700 to 900 | 1680-2150 | 15 to 25 | < 10 | Jørgensen (1991); Oldham and Gerhardt (1975) |
| *Hyla versicolor* | 1100 | 1300 to 1800 | 1300 to 1600 | 2200 | 15 to 25 | < 6 | Jørgensen and Gerhardt (1991) |
| *Hyla chrysoscelis* | 1250 | 1600 to 1900 | 1400 | 2500 | 10 to 15 | < 10 | Caldwell et al. (2014); (Ward et al., 2013) |
| *Eleutherodactylus coqui* | 1500 | 1720 | 1500 to 1800 | 2400-2700 | 10 to 15 | 5 to 6 | Jørgensen et al. (1991); (Narins and Smith, 1986); (Lopez and Narins, 1991) |
| *Rana temporaria* | 350 to 500 | 600 to 1000 | 650 to 700 | 1200 | 10 to 15 | < 5 | Jørgensen (1991); (Brzoska et al., 1977) |

Our findings extend this earlier work by showing that sound transmission through the lungs does not improve directional hearing and thus does not function in localizing conspecific calls. This conclusion is at odds with early suggestions that the lungs might improve localization of conspecific calls (Ehret et al., 1990) and with previous studies suggesting that sound transmission through the lungs improves the tympanum’s directionality (Jørgensen, 1991; Jørgensen et al., 1991). Previous studies demonstrating an impact of the lungs on directional hearing have inferred changes in VADs in small numbers of subjects (e.g., *N* ≤ 6) after applying thick layers of petroleum jelly to the body wall to dampen sound transmission to inflated lungs (Jørgensen, 1991; Jørgensen et al., 1991) or after directly manipulating the state of lung inflation (Caldwell et al., 2014; Jørgensen, 1991). In their study of the coqui frog, for example, Jørgensen et al. (1991) reported that dampening sound transmission through the body wall with petroleum jelly eliminated the dip in the tympanum’s transfer function and decreased the tympanum’s vibration amplitude by 10-15 dB at low frequencies typical of the “co” note of the advertisement call. Interestingly, they also report that the maximum directionality (inferred from the VAD) was “only slightly reduced” in treated frogs (p. 228, Jørgensen et al., 1991). In green treefrogs, deflating the lungs decreased the maximum VAD by 4 dB, from 15.5 dB to 11.5 dB (Table 1). Thus, while the lungs influenced the magnitude of the VAD at frequencies near the lung resonance, a large directionality remained in the deflated state, presumably due to the internal coupling of the tympana. These data suggest both a relatively weak influence of lung inflation on interaural transmission gain and that the lungs do not contribute much to interaural coupling and hence to directionality. One important consideration for interpreting these findings based on the VAD is that the VAD measures the directional response of a single tympanum, whereas frogs must use binaural comparisons to localize conspecific calls in azimuth (Feng et al., 1976; Rheinlaender et al., 1979). Hence, IVADs are the more relevant measure of directional hearing because they characterize the binaural disparities in tympanum vibrations that are ultimately used by the nervous system to localize sound sources (Jørgensen et al., 1991). The IVAD can even be seen as a simplified model of interaural comparison in the CNS, assuming that the periphery is symmetrical across the midline (Christensen-Dalsgaard and Manley, 2005). The magnitude of IVADs reported here (e.g., 4 dB to 10 dB) are in line with those reported in earlier studies of coqui frogs (8 dB; Jørgensen et al., 1991), eastern gray treefrogs (e.g., 3 dB to 7 dB; Jørgensen and Gerhardt, 1991), and Cope’s gray treefrogs (e.g., 2 dB to 5 dB; Caldwell et al., 2014). Like VADs, IVADs tend to be larger at frequencies near the lung resonance and the dip in the tympanum’s transfer function and smaller at frequencies emphasized in advertisement calls (Bee and Christensen-Dalsgaard, 2016). However, despite larger IVADs at frequencies intermediate between call frequencies, sound localization performance is actually degraded when signals are modified to emphasize the frequencies of greatest directionality (Jørgensen and Gerhardt, 1991). No previous study has investigated the potential effects the lungs may have on the magnitude of the IVAD. Our data indicate unequivocally that the state of lung inflation has negligible impact on IVADs: differences in IVADs between the inflated and deflated conditions were close to 0 dB across frequency and azimuth. Because IVADs are likely the most important cue for sound localization in frogs, these findings suggest the auditory nervous system receives consistent directional information from the periphery, probably based on the interaural coupling of the tympana, irrespective of the state of lung inflation.

To date, no study has investigated the influence of the lungs on phase-related cues for sound localization, such as the magnitude of interaural phase differences (i.e., interaural vibration phase difference, or IVPD) and the corresponding interaural time differences (i.e., interaural vibration time difference, or IVTD), which might provide cues for call localization in some species, particularly for low-frequency spectral components (Ho and Narins, 2006) or amplitude-modulated signals (Klump et al., 2004). It seems unlikely that phase differences in the spectral components of calls would be useful for call localization in green treefrogs. In their study of the congeneric Cope’s gray treefrog (*H. chrysoscelis*), Caldwell et al. (2014) reported that IVPDs were generally similar to or smaller than phase differences at the external surfaces of the two tympana. In that study, IVTDs in frogs with inflated lungs never exceeded 120 μs and most were less than 60 μs, even for frequencies as low as 600 Hz. Moreover, as previously noted by Klump et al. (2004), phase locking in the frog auditory nerve is weak at frequencies near the low-frequency component of green treefrogs calls and does not occur at frequencies near the high-frequency component of their calls (Feng et al., 1991; Narins and Hillery, 1938; Narins and Wagner, 1989; Ronken, 1990; Rose and Capranica, 1985). Thus, any influence the lungs may have on the magnitude of IVPDs would likely be inconsequential for directional hearing based on the use of interaural phase cues.

The mismatch between the frequencies of maximal tympanum directionality and the frequencies emphasized in conspecific calls has remained paradoxical: why is the tympanum most directional at frequencies not used for communication given the importance of sound localization in frog sexual and social behavior? Several hypotheses have been proposed to explain this apparent paradox. Jørgensen et al. (1991) hypothesized that in coqui frogs (*E. coqui*) individuals might be able to tune the maximal directionality of the tympana to the frequency of the “co” note of the advertisement call by adjusting the volume, and hence resonance frequency, of their lungs. While the frequency of maximal directionality can be altered by changing the lung’s volume (Jørgensen, 1991), current evidence suggests changes in lung inflation produce relatively small changes. For example, reducing the lung volume from 100% to 30% of maximal inflation only changed the lung resonance frequency by 150 Hz in barking treefrogs (*H. gratiosa*) (Jørgensen, 1991). In the present study, deflating the lungs shifted the frequency of the maximum VAD downward by only 231 Hz (from 1486 Hz to 1255 Hz; Table 1) and the frequency of the maximum IVAD downward by 300 Hz (from 1550 Hz to 1250 Hz; Table 2). Both shifts were insufficient to reach the lower spectral peak of the advertisement call at 834 Hz. These findings suggest the scope for tuning the tympanum’s directionality by manipulating lung resonance via volume changes is rather limited. Additional hypotheses for the documented mismatch are that it reflects either an evolutionary adaptation to avoid overlap between call frequencies and the frequencies affected by the lung input because of variability of the lung-induced directionality during the respiratory cycle or a physical constraint between lung resonance frequency and call frequencies (Jørgensen, 1991). Both hypotheses are unsatisfactory. As already noted, even large changes in lung inflation produce only small changes in the frequency of maximal directionality. Moreover, the frog’s lungs remain continuously pressurized above atmospheric pressure during the respiratory cycle (De Jongh and Gans, 1969), and they remain inflated for relatively long periods punctuated by brief episodes of ventilation when pulmonary air is expelled and then refilled using an active pump mechanism driven by buccal musculature (Jørgensen et al., 1991). Thus, the acoustical properties of the lung input probably change very little over the respiratory cycle, and frogs probably do not contend with large variability in directionality due to pulmonary respiration. There is also no clear co-dependence between call frequencies and lung resonance frequency, since call frequencies depend on the size and structure of vocal cords and cricoid cartilages, and the lung resonance frequency, measured in the inflated, static situation, is not really relevant in call production, during which the lungs deflate.

More recently, Lee et al. (2020) proposed a resolution to the paradoxical mismatch between call frequencies and lung-mediated effects on the frequencies of maximal directionality. They provided evidence from green treefrogs suggesting it is actually the lung-mediated reduction in sensitivity associated with the dip in the tympanum’s transfer function, and not enhanced directionality, that is adaptive in the context of communication and environmental noise control. By comparing transfer functions with the lungs inflated versus deflated, they found that the lung resonance creates a directionally tuned notch filter that attenuates the tympanum’s vibration amplitude in the frequency range of 1400 Hz to 2200 Hz and predominantly within the contralateral portion of the frontal hemifield. A physiological model of peripheral frequency tuning indicated that these lung-mediated reductions in tympanum sensitivity are restricted to a frequency range where the tuning of the frog’s two inner ear organs for transducing airborne sound frequencies, the amphibian and basilar papillae, overlaps at the high sound levels used for communication. Based on these findings, Lee et al. (2020) suggested the lung-to-ear sound transmission pathway functions to sharpen peripheral frequency tuning. They argued that inflated lungs create a mechanism for real-time spectral contrast enhancement (SCE) similar to that implemented in signal processing algorithms in hearing aids and cochlear implants that can improve speech recognition in noise by humans with impaired hearing (Baer et al., 1993; Nogueira et al., 2016; Simpson et al., 1990). For frogs, the end result of spectral contrast enhancement would be a lung-mediated improvement in peripheral matched filtering in the spectral domain, which is believed to play important roles in call recognition as a mechanism to reduce interference from heterospecific signals or environmental noise (Simmons, 2013). To evaluate this possibility, Lee et al. (2020) integrated social network analyses of content-scale citizen science data on frog calling activity with bioacoustic analyses of calls to show that some of the most common species encountered in mixed-species choruses across the green treefrog’s geographic range produce calls with prominent spectral peaks in the range of 1400 Hz to 2200 Hz. Hence, the lung-to-ear sound transmission pathway may serve an important noise control function in frogs that communicate in mixed species choruses. By showing that the lungs do not improve directional hearing in the context of localizing conspecific calls, the current study suggests alternative adaptive functions of the lung-to-ear sound transmission pathway, such as spectral contrast enhancement (Lee et al., 2020) or hearing protection during calling (Narins, 2016), deserve additional consideration.

## Acknowledgements

We thank Chris Maldonado of the Texas Parks and Wildlife Division and Gary Calkins of the East Texas Conservation Center for permission to collect frogs under Scientific Permit Number SPR-0410-054; J. Tanner and M. Elson for help collecting frogs; and S. Gupta for assistance with animal husbandry.

## Author contributions

J.C.D., N.L., and M.A.B. designed the experiments, collected and analyzed the data, and co-wrote the manuscript.

